## Supplemental information for "A method to estimate absolute odorant concentration of olfactory stimuli"

#### Model of odorant release from a liquid to a headspace compartment

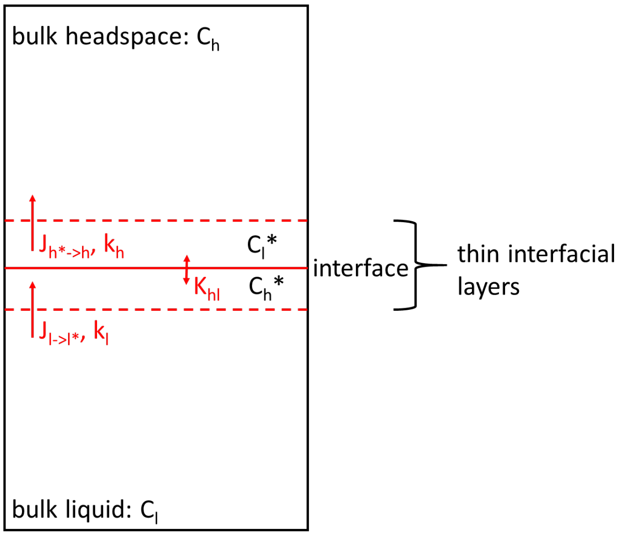


We consider a system composed of a liquid and a headspace compartment, separated by an interface (the liquid's surface).

Hypothesis 1: The bulk liquid and bulk headspace compartments are perfectly stirred, i.e. odorant concentration is always uniform across their whole volume: noted $C_l$ and $C_h$. This is at the exception of two thin boundary layers on each side of the interface.

Hypothesis 2: Transport across the interface itself is instantaneous, so the two interfacial layers are always at thermodynamic equilibrium with one another. At any moment:

$$C_h^* = K_{hl} * C_l^* \quad \text{(A1)}$$

Mass fluxes from bulk liquid to liquid interfacial layer $J_{l\text{->}l^*}$ and from headspace interfacial layer to bulk headspace $J_{h^*\to h}$ (mol m$^{-2}$ s$^{-1}$) depend on the difference between bulk and interfacial concentrations and on transfer coefficients $k_l$ and $k_h$ (m/s):

$$J_{l\to l*} = k_l * (C_l - C_l^*) \quad \text{(A2)}$$

$$J_{h^*\to h} = k_h * (C_h^* - C_h) \quad \text{(A3)}$$

If multiplied by $K_{hl}/k_l$, equation (A2) becomes

$$J_{l\to l*} * \frac{K_{hl}}{k_l} = K_{hl} * (C_l - C_l^*) \quad \text{(A4)}$$

When incorporating equation (A1) and dividing by $k_h$, equation (A3) becomes

$$J_{h*\to h} * \frac{1}{k_h} = K_{hl} * C_l^* - C_h \quad \text{(A5)}$$

Hypothesis 3: No accumulation occurs within the interfacial layers. The mass flow into the liquid interfacial layer is equal to the mass flow out from the headspace interfacial layer, and equal to $J_{l\text{->}h}$ the global mass flow through the interface from bulk liquid to bulk headspace:

$$J_{l\to l*} = J_{h*\to h} = J_{l\to h} \quad \text{(A6)}$$

This allows to deduce the value of $J_{l\text{->}h}$ : summing up equations (A4) and (A5) and incorporating equation (A6) we get

$$J_{l\to h} * \frac{K_{hl}}{k_l} + J_{l\to h} * \frac{1}{k_l} = K_{hl} * (C_l - C_l^*) + K_{hl} * C_l^* - C_h$$

which simplifies to

$$J_{l\to h} * \left(\frac{1}{k_h} + \frac{K_{hl}}{k_l}\right) = K_{hl} * C_l - C_h$$

and further to

$$J_{l\to h} = k_{glob} * (K_{hl} * C_l - C_h) \quad \text{(A7)}$$

where $k_{glob}$ is the global mass transfer coefficient, defined as

$$\frac{1}{k_{glob}} = \frac{1}{k_h} + \frac{K_{hl}}{k_l} \quad \text{(A8)}$$

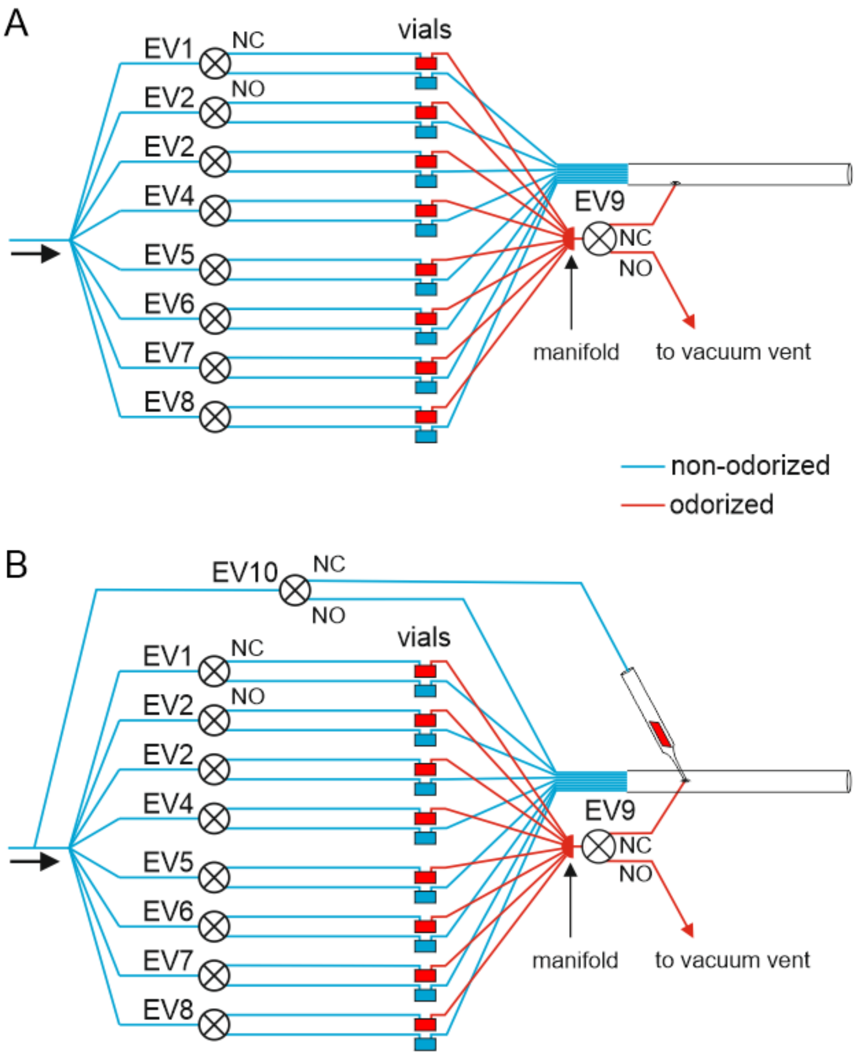


**Figure S1 -** Odor delivery devices used for EAG (A) and SSR recordings (B). Compared to the device presented in Figure 1A and characterized in this work, it included two additional electrovalves (EV9 and EV10) but the length of Teflon tubing was kept similar. For EAG and SSR, EV9 allowed delivering a constant VPC concentration after the elimination of the initial concentration peak. VPC source outlets merge into a low dead volume manifold (MPP8, Warner Instruments, Holliston, MA, USA) connected to EV9. When EV9 was closed, air was directed to vacuum vent and when it was open, air was directed to the side hole of the glass tube via a 1-cm Teflon tubing. Non-odorized vial outlets remained connected to the glass tube and provided the carrier airflow. For SSR, EV10 was used to deliver pheromone pulses on a background of VPC.

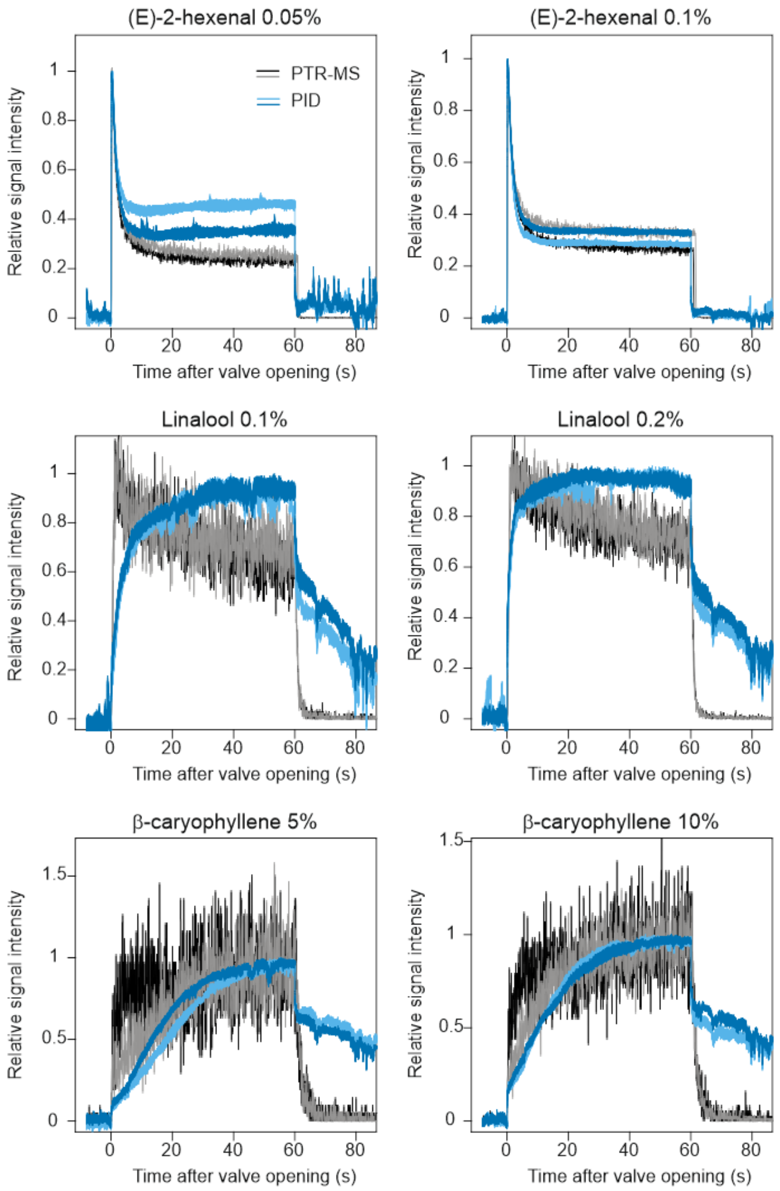


**Figure S2 -** Comparison of stimulus time courses at the outlet of the source, as observed by a photoionisation detector (PID) and a proton-transfer reaction mass spectrometer (PTR-MS). The identity and concentration of odorant loaded inside the source is indicated above each panel.

**Table S1 - Literature review of ambient air concentrations of Green Leaf Volatiles (GLVs) in natural/agricultural landscapes.**

Publications where authors have tentatively or conclusively identified GLVs in the ambient air of the studied ecosystem, within 35 m above the vegetation canopy. We extracted the estimates of average, median, maximum or range of GLV concentrations. Note that the reported values always represent an average over a certain volume of sampled air. The PTR-MS detects and quantifies the products of a gas sample's ionization, classified by their mass to charge ratio (m/z). When a complex sample such as ambient air is analyzed, it is not possible to discriminate among individual compounds with the same m/z. Identification of atmosphere components by PTR-MS therefore remain tentative unless further analyses confirm which VPCs actually contribute to the detected ions. Ionization of hexenyl acetate produces notably m/z 83 (abundant but unspecific) and m/z 143 (specific but much less abundant [1]). Several other GLVs generate mostly m/z 83 when ionized. This table summarizes the reported concentrations of (**A**) PTR-MS ions to which Z3HA, (**B**) PTR-MS ions that are markers of other GLVs, and (**C**) GLV concentrations estimated by other methods. LOD = limit of detection, sd = standard deviation, ppbv = parts per billion by volume.

**Table S1A PTR-MS data: m/z 83 and 143**

| Reference | Ecological context | ion | author's identification | Reported concentrations (ppbv) | | notes |
|---|---|---|---|---|---|---|
| [2] | Creosote (*Larrea tridentata*) bushland, Arizona summer, 3m above canopy | m/z 83 | 3-methyl furan, hexanal and 3-hexenyl acetate | 0.1-0.15 | range, noon time | Contributor compounds confirmed by GC-PTR-MS and GC-MS on cresoste branch headspace extracts |
| | | | | 0.01-0.1 | range, 48h time series | |
| | | m/z 143 | nonanal, 3-hexenyl acetate, 1-chloro-2-methoxy benzene | 0.6-1 | range, noon time | |
| | | | | 0-0.6 | range, 48h time series | |
| [3] | *Pseudotsuga menziesii* forest, Denmark summer, under, in and up to 13m above canopy | m/z 83 | hexenols, hexanal (E,Z)-2-hexenyl acetate | 0,4 | above canopy average | LOD 0.6-0.7 ppbv |
| | | | | 0.18-0.34 | averages per time of day and height in canopy | |
| [4,5] | *Citrus sinensis* orchard, California year round, in and up to 5m above canopy | m/z 83 | hexanal, hexenols | 0.14, 0.25, 0.15 | midday averages during winter, flowering and summer | LOD 0.02 ppbv occasional peaks up to 4ppbv during flowering |
| | | m/z 83.086* | hexanal, hexenols | 0.07, 0.17 | day and night time averages in summer | m/z 143.114 (=hexenyl acetate) not in the list of detected ions |
| [6] | Palm oil plantation, Borneo april-july 3m above canopy | m/z 83 | hexanal | 0,5 | global average | |
| | | | | 1 ± 0.5 | midday average ± sd | |
| [6] | Rainforest Borneo april to july 35 m above canopy | m/z 83 | hexanal | very low | | |
| [7] | Rainforest Borneo april to july, 35 m above canopy | m/z 83 | hexanal, hexenols | 0.06±0.02 | global mean and variance | LOD 0.04 ppbv |
| | | | | 0.06, 0.04-0.09 | midday average and range | |
| | | | | 0.04, 0.02-0.06 | night time average and range | |
| [8] | Permanent managed grassland, Swizerland summer 1.2m above ground during and after grass cutting/hay removal | m/z 83 | (Z)-3-hexenol, (E)-3-hexenol, (E)-2-hexenol, hexanal, (E,Z)-3-hexenyl acetate | 0.92, 0.19, 0.22, 0.05 | day time average, 1 day, 2 days, 3 days and 4 days after grass cutting | day 4 concentration comparable to pre-cutting (Z)-3-hexenyl acetate represents 60-90% of m/z 83 from dominant grass species emissions (GC-PTR-MS on headspace) |
| [9] | mature rainforest, Amazonia during a drought event, under, in and up to 10 m above canopy | m/z 83 | 3-methyl furan, hexenols | 0.05-0.5, 0.2-0.8 | monthly averages, mid-canopy and above canopy | Identity of contributor compounds confirmed by GC-PTR-MS on above canopy air. Undetectable at ground level |
| [10] | Urban lawn, Austria before, during and after mowing | m/z 83 | hexanal, hexenols | up to 2-6 | peaks after mower passage | Very low concentration before mowing m/z 143 not detected |
| | | | | 0-0.7 | 3 to 8h post mowing | |
| [11] | Oak-dominated forest in agricultural landscape, Italy summer, 4m above canopy | m/z 83.086* | fragment of C6 GLVs | 0.8 | whole campaign average | m/z 101.0961 (parent ion for hexanal and hexenols) is detected. m/z 143.114 (parent ion for hexenyl acetate) is not. |

**Table S1B PTR-MS data: other GLV characteristic ions**

| Reference | Ecological context | ion | Author's identification | Concentrations (ppbv) | | notes |
|---|---|---|---|---|---|---|
| [4] | Citrus sinensis orchard, California<br>year round, in and up to 5m above canopy | m/z 99 | hexenal | 0.03, 0.04, 0.06 | midday averages during winter, flowering and summer | LOD 0.14 ppbv<br>occasional peaks up to 4ppbv during flowering season |
| [5] | | m/z 99.078* | hexenal | 0.04, 0.05 | day and light time averages in summer | |
| [6] | Rainforest Borneo<br>april to july, 35 m above canopy | m/z 85 | hexenols | very low | | |
| | Palm oil plantation, Borneo<br>3 m above canopy | m/z 85 | hexenols | very low | | |
| [8] | Permanent managed grassland on Swiss plateau<br>summer 1.2m above ground<br>during and after grass cutting/hay removal | m/z 81 | (E)-2-hexenal, (Z)-3-hexenal, pinene fragments | 0.82, 0.16, 0.17, 0.05 | day time average, 1 day, 2 days, 3 days and 4 days after grass cutting | day 4 concentration comparable to pre-cutting<br>(Z)-3-hexenal represents 100% of m/z 81 from most dominant grass species emissions (GC-PTR-MS on headspace) |
| [9] | mature rainforest, Amazonia<br>during a drought event, under, in and up to 10m above canopy | m/z 85 | hexanol | 0.04-0.16 | monthly average mid canopy and above canopy | Undetectable at ground level.<br>GC-PTR-MS analysis of above canopy air reveals only 1 peak |
| [10] | Urban lawn, Austria<br>before, during and after mowing | m/z 81 | hexenal | <0.1 | before mowing | |
| | | | | 2-6 | peaks post mower passage | |
| | | | | 0-0.7 | 3 to 8h post mowing | |
| [12,13] | Boreal forest dominated by *Pinus sylvestris*, Finland<br>year round, in canopy | m/z 101 | hexanal, (Z)-3-hexenol | 0.22-2.11 | monthly medians | LOD 0.22 |
| | | m/z 99 | hexenals | 0.05 | global median | |

**Table S1C GLV concentrations estimated by other methods**

| Reference | Ecological context | technique | Compounds | Concentrations (ppbv) | |
|---|---|---|---|---|---|
| [14,15] | Spruce (*Picea abies*) forest, Bavaria summer, in and 5 m above canopy | HPLC | trans-2-hexenal | 0.05-0.55 | whole campaign ranges |
| | | | nonanal | 0-0.45 | |
| | | | hexanal | 0-0.28 | |
| | | | heptanal | 0-0.15 | |
| | | | octanal | 0.0.25 | |
| | | | decanal | 0-0.02 | |
| [16] | Hemiboreal forest, dominated by *Picea abies*, Estonia year round, under and in canopy by clear sky weather only | GC-MS, headspace | (Z)-3-hexenol | 0.001-1 | averages per month and height |
| | | | 1-hexanol | 0.0005-1.3 | averages per month and height |

* measured by PTR-ToF-MS. This instrument measures mass to charge ratio with enough precision to deduce the molecular formula of the detected ion.

### References


[1] Fall R (1999) Volatile organic compounds emitted after leaf wounding: On-line analysis by proton-transfer-reaction mass spectrometry. *J Geophys Res Atm* 104(D13):15,963-15,974. DOI: 10.1029/1999JD900144

[2] Jardine K, Abrell L, Kurc SA, Huxman T, Ortega J, Guenther A (2010) Volatile organic compound emissions from *Larrea tridentata* (creosotebush). *Atmos. Chem. Phys.* 10:12191–12206. DOI: 10.5194/acp-10-12191-2010

[3] Copeland N, Cape JN, Nemitz E, Heal MR (2014) Volatile organic compound speciation above and within a Douglas fir forest. *Atmos Environ* 94:86-95. DOI: 10.1016/j.atmosenv.2014.04.035

[4] Fares S, Park JH, Gentner DR, Weber R, Ormeño E, Karlik J, Goldstein AH (2012) Seasonal cycles of biogenic volatile organic compound fluxes and concentrations in a California *citrus* orchard. *Atmos. Chem. Phys.* 12:9865–9880. DOI: 10.5194/acp-12-9865-2012

[5] Park JH, Goldstein AH, Timkovsky J, Fares S, Weber R, Karlik J, Holzinger R (2013) Eddy covariance emission and deposition flux measurements using proton transfer reaction – time of flight – mass spectrometry (PTR-TOF-MS): comparison with PTR-MS measured vertical gradients and fluxes. *Atmos. Chem. Phys.* 13:1439–1456. DOI: 10.5194/acp-13-1439-2013

[6] MacKenzie AR, Langford B, Pugh TA, Robinson N, Misztal PK, Heard DE, Lee JD, Lewis AC, Jones CE, Hopkins JR, Phillips G, Monks PS, Karunaharan A, Hornsby KE, Nicolas-Perea V, Coe H, Gabey AM, Gallagher MW, Whalley LK, Edwards PM, Evans MJ, Stone D, Ingham T, Commane R, Furneaux KL, McQuaid JB, Nemitz E, Seng YK, Fowler D, Pyle JA, Hewitt CN (2011) The atmospheric chemistry of trace gases and particulate matter emitted by different land uses in Borneo. *Philos Trans R Soc Lond B Biol Sci* 366(1582):3177-3195. DOI: 10.1098/rstb.2011.0053

[7] Langford B, Misztal PK, Nemitz E, Davison B, Helfter C, Pugh TAM, MacKenzie AR, Lim SF, Hewitt CN (2010) Fluxes and concentrations of volatile organic compounds from a South-East Asian tropical rainforest. *Atmos. Chem. Phys.* 10(17):8391-8412. DOI: 10.5194/acp-10-8391-2010

[8] Davison B, Brunner A, Ammann C, Spirig C, Jocher M, Neftel A (2007) Cut-induced VOC emissions from agricultural grasslands. *Plant Biol* 9:e60-e68. DOI: 10.1055/zs-2007-965043

[9] Jardine KJ, Chambers JQ, Holm J, Jardine AB, Fontes CG, Zorzanelli RF, Meyers KT, de Souza VF, Garcia S, Gimenez BO, Piva LR, Higuchi N, Artaxo P, Martin S, Manzi AO (2015) Green leaf volatile emissions during high temperature and drought stress in a central Amazon rainforest. *Plants* 4(3):678-690. DOI: 10.3390/plants4030678

[10] Karl T, Fall R, Jordan A, Lindinger W (2001) On-line analysis of reactive VOCs from urban lawn mowing. *Environ. Sci. Technol.* 35(14):2926-2931. DOI: 10.1021/es010637y

[11] Schallhart S, Rantala P, Nemitz E, Taipale D, Tillmann R, Mentel TF, Loubet B, Gerosa G, Finco A, Rinne J, Ruuskanen TM (2016) Characterization of total ecosystem-scale biogenic VOC exchange at a Mediterranean oak–hornbeam forest. *Atmos. Chem. Phys.* 16(11):7171-7194. DOI: 10.5194/acp-16-7171-2016

[12] Ruuskanen TM, Taipale R, Rinne J, Kajos MK, Hakola H, Kulmala M (2009) Quantitative long-term measurements of VOC concentrations by PTR-MS: annual cycle at a boreal forest site. *Atmos. Chem. Phys. Discuss.* 9(1):81-134. DOI: 10.5194/acpd-9-81-2009

[13] Patokoski J, Ruuskanen TM, Hellen H, Taipale R, Gronholm T, Kajos MK, Petaja T, Hakola H, Kulmala M, Rinne J (2014) Winter to spring transition and diurnal variation of VOCs in Finland at an urban background site and a rural site. *Boreal Environ. Res.* 19(2):79-103. DOI: 10138/165174

[14] Klemm O, Held A, Forkel R, Gasche R, Kanter H-J, Rappenglück B, Steinbrecher R, Müller K, Plewka A, Cojocariu C, Kreuzwieser J, Valverde-Canossa J, Schuster G, Moortgat GK, Graus M, Hansel A (2006) Experiments on forest/atmosphere exchange: Climatology and fluxes during

two summer campaigns in NE Bavaria. *Atmos. Environ.* 40:3-20. DOI: 10.1016/j.atmosenv.2006.01.060

[15] Müller K, Haferkorn S, Grabmer W, Wisthaler A, Hansel A, Kreuzwieser J, Cojocariu C, Rennenberg H, Herrmann H (2006) Biogenic carbonyl compounds within and above a coniferous forest in Germany. *Atmos. Environ.* 40:81-91. DOI: 10.1016/j.atmosenv.2005.10.070

[16] Noe SM, Hüve K, Niinemets Ü, Copolovici L (2012) Seasonal variation in vertical volatile compounds air concentrations within a remote hemiboreal mixed forest. *Atmos. Chem. Phys.* 12(9):3909-3926. DOI: 10.5194/acp-12-3909-2012
